# Efficient CRISPR/Cas12a-based genome editing toolbox for metabolic engineering in *Methanococcus maripaludis*

**DOI:** 10.1101/2021.12.29.474413

**Authors:** Jichen Bao, Enrique de Dios Mateos, Silvan Scheller

## Abstract

The rapid-growing and genetically tractable methanogen *Methanococcus maripaludis* is a promising host organism for the biotechnological conversion of carbon dioxide and renewable hydrogen to fuels and value-added products. Expansion of its product scope through metabolic engineering necessitates reliable and efficient genetic tools, particularly for genome edits to the primary metabolism that affect cell growth. Here, we have designed a genome editing toolbox by utilizing Cas12a from *Lachnospiraceae bacterium* ND2006 (LbCas12a) in combination with the homology-directed repair machinery endogenously present in *M. maripaludis*. Remarkably, this toolbox can knock out target genes with a success rate of up to 95%, despite the hyper-polyploidy of *M. maripaludis*. For the purposes of demonstrating a large-sized deletion, we have replaced the flagellum operon (ca. 8.9 kbp) by the β-glucuronidase gene. To facilitate metabolic engineering and flux balancing in *M. maripaludis*, the relative strength of 15 different promoters were quantified in the presence of the two common growth substrates, formate or carbon dioxide and hydrogen. This CRISPR/LbCas12a toolbox can be regarded as a reliable and fast method for genome editing in a methanogen.

## INTRODUCTION

Methanogenic archaea are biotechnologically employed in a variety of uses, e.g. for methane production in anaerobic digestors^1^, as biocatalysts in power-to-gas processes^2^, and as promising host organisms for synthetic pathways to convert carbon dioxide (CO_2_) into value-added products^3,4^. Hydrogenotrophic methanogens utilize the reductive acetyl-CoA pathway for CO_2_ fixation^5^, an energy-efficient route to synthesize organic carbon from CO_2_ and hydrogen (H_2_), which is similar to that found in acetogens^6^. However, subtle differences exist between the acetogenic^7^ and methanogenic pathways of CO_2_ reduction in terms of ATP investment and cofactor utilization^8^. Depending on the requested product that needs to be generated from CO_2_ as the carbon source, methanogens may be better suited host organisms than acetogens. Recently, the methanogen *Methanosarcina acetivorans* was re-engineered to no longer depend on methane production for its energy metabolism^9^, and thereby it might serve as an example where a methanogen could be utilized for generating an increased repertoire of new potential products, besides just methane.

*Methanococcus maripaludis* is a promising methanogenic host organism for metabolic engineering of CO_2_-fixation pathways due to its advantageous growth properties, e.g., two-hour doubling time^10,11^, moderate growth temperature of 38°C, and ability to fix nitrogen^12,13,14^. Typical electron donors for CO_2_ reduction in *M. maripaludis* include formate, H_2_, and bioelectrically coupled systems^15,16^. Efforts to expand the product scope beyond methane for *M. maripaludis* are already underway. As an example, the mevalonate pathway in this methanogen was metabolically engineered to produce geraniol from CO_2_ and formate^4^. Even so, efficient and reliable genome editing tools are critical for successful metabolic engineering in *M. maripaludis*. Marker recycling is prerequisite for multi target engineering. In the case of *M. maripaludis*, although a ‘pop-in/pop-out’ markerless-based genome editing technique has been developed^17^, it tends to have a problematic low positive rate, which can sometimes be less than 5%^17^, especially for those modifications that affect cell growth. As an alternative, the CRISPR/Cas (clustered regularly interspaced short palindromic repeats/CRISPR associated protein) system might remedy this problem because of its reputation for highly efficient genome editing.

The CRISPR/Cas9 system has already been successfully used for genome editing in a variety of organisms^18,19,20,21,22,23^ owing to its simplicity and high efficiency. However, only a few CRISPR genome editing toolboxes have been developed for archaea^24^. The first CRISPR/Cas9-mediated genome editing system for a methanogen was reported in 2017 using *M. acetivorans* as the model organism^25^. This Cas9-based system recognizes and cleaves a 20 nt target sequence that is flanked by a 3’-NGG protospacer adjacent motif (PAM). This contrasts with Cas12a, which instead recognizes the 5’-thymine (T)-rich PAM 5’-TTTV. This recognition site makes Cas12a the better option for developing a CRISPR toolbox in microbes with an adenine (A)- and T-rich genome. Another advantageous attribute of Cas12a lies in its ribonuclease activity, which allows the formation of multiple guide RNAs (gRNAs) from a single transcript^26,27^. Since the *M. maripaludis* genome has a high AT content (67.1%), we decided to use the Cas12a from *Lachnospiraceae bacterium* ND2006 (LbCas12a) and combine it with the intrinsic homology-directed repair machinery to develop a CRISPR genome editing toolbox. In our study, we examined how the length of the repair fragment (RF) and the distance of the RF to the double stranded break (DSB) can impact on the genome-editing efficiency. To further expand the versatility and editing potential of this genetic toolbox, we also established a Cas9-based editing system. As an application of our toolbox, we deleted the *M. maripaludis* flagellum operon and replaced it by the β-glucuronidase gene. Finally, we were also able to quantify the relative strength of 15 different *M. maripaludis* promoters in the presence of the two common growth substrates formate or H_2_+CO_2_.

## RESULTS AND DISCUSSION

### CRISPR/Cas12a-based introduction of double-stranded breaks and transformation efficiency

The host strain *M. maripaludis* JJΔupt was used for all experiments, applying the natural transformation protocol recently established^28^. The *Escherichia coli/M. maripaludis* shuttle vector pLW40^29^ was used as a backbone to construct the final toolbox plasmid pMM002P (Table 1). The schematic diagram of the toolbox plasmid and description of its components is given in Fig.1a. The transformation efficiency of pMM002P in JJ△upt was ca. 5 × 10^4^ colony forming units per 2 *μg* DNA [cfu (2 *μg* DNA)^-1^](Fig. 1b), which is higher than the reported efficiency of the empty plasmid pLW40neo [2.4 × 10^3^ cfu (2 *μ*g DNA)^-1^]^28^. The high transformation efficiency indicates that expressing LbCas12a alone is not toxic to the host cell. When one gRNA was co-expressed with LbCas12a, only 3-18 colonies could be observed on the transformation plates. When two gRNAs were co-expressed, only 0-3 colonies were formed (Fig. 1b). These results indicated that the LbCas12a-gRNA complex can perform a lethal double stranded break in *M. maripaludis*. Nonhomologous end joining (NHEJ) machineries are rare in archaea^30^, and we could not find the homologous Ku protein in *M. maripaludis* JJ. Thus, the NHEJ does not cause the escape from the cleavage.

**Figure 1.**
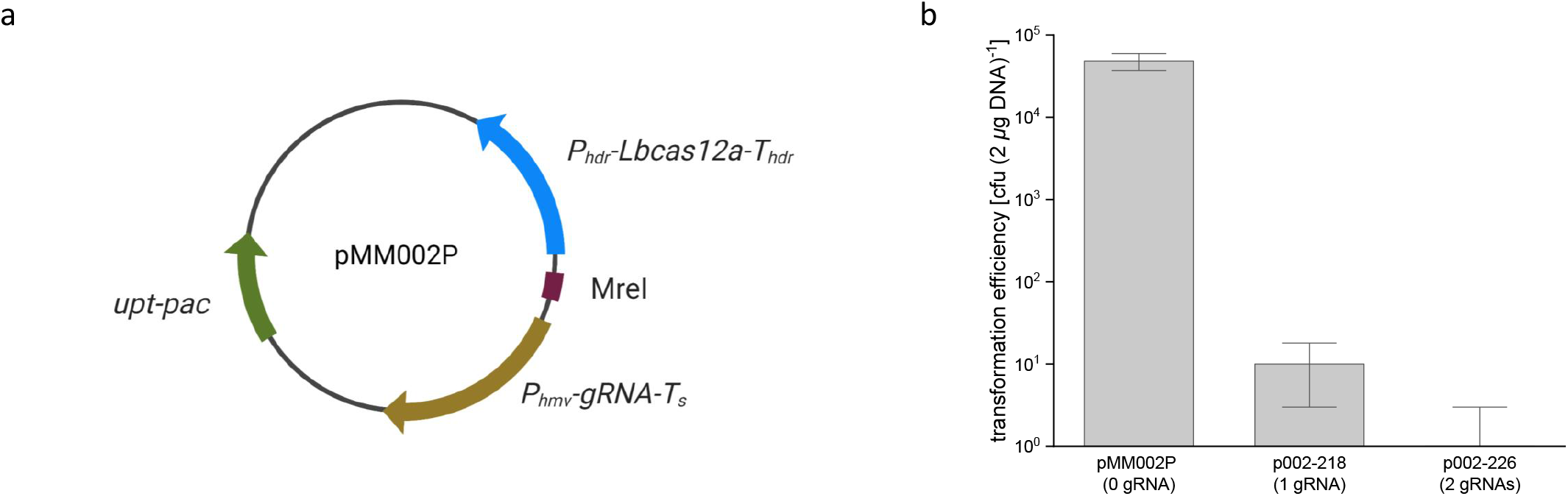
Features and transformation efficiency of the CRISPR/LbCas12a plasmid pMM002P. a) The schematic diagram of pMM002P. The uracil phosphoribosyltransferase (*upt*) from *M. maripaludis* S2 as a counter-selective marker is co-expressed with the codon-optimized puromycin *N*-acetyltransferase (*pac*) under the *P_mcr_* promoter^4^. LbCas12a expression is driven by the promoter *P_hdr_* from *Methanococcus voltae* A3. The histone promoter *P_hmv_* from *M. voltae* A3 is used to express the gRNA. For inserting the required gRNA, two *PaqC* I sites are placed between the direct repeat sequence and the synthetic terminator in the opposite direction for spacer fusion (not displayed). The gRNA of the plasmid pMM002P containing two *PaqC* I sites is designed to not target the native chromosome. An *Mre* I restriction site was assigned between the promoters *P_hdr_* and *P_hmv_* for RF insertion, if needed. b) Cleavage test with transformed plasmid pMM002P, using 0, 1, or 2 gRNA carried out in triplicates. The error bars represent the standard deviation (n=3).

### Genome editing via CRISPR/LbCas12a by providing a repair fragment on the same plasmid

A gRNA was expressed on pMM002P (p002-218) targeted to the gene *flaI* within the flagellum operon. The lethal effect of the functional gRNA could be relieved by including RFs with various lengths of the homology arms (Fig. 2a). When a 1000 bp long homologous arm on each side was provided, the transformation efficiency reached 2.8 × 10^4^ cfu (2 *μ*g DNA^-1^). We therefore used the 1000 bp long homologous arms as the standard condition in the following experiments. 250 bp long homologous arms would be enough to rescue the DSB generated by Cas12a/gRNA complex, but the transformation efficiency is 70 times lower.

**Figure 2.**
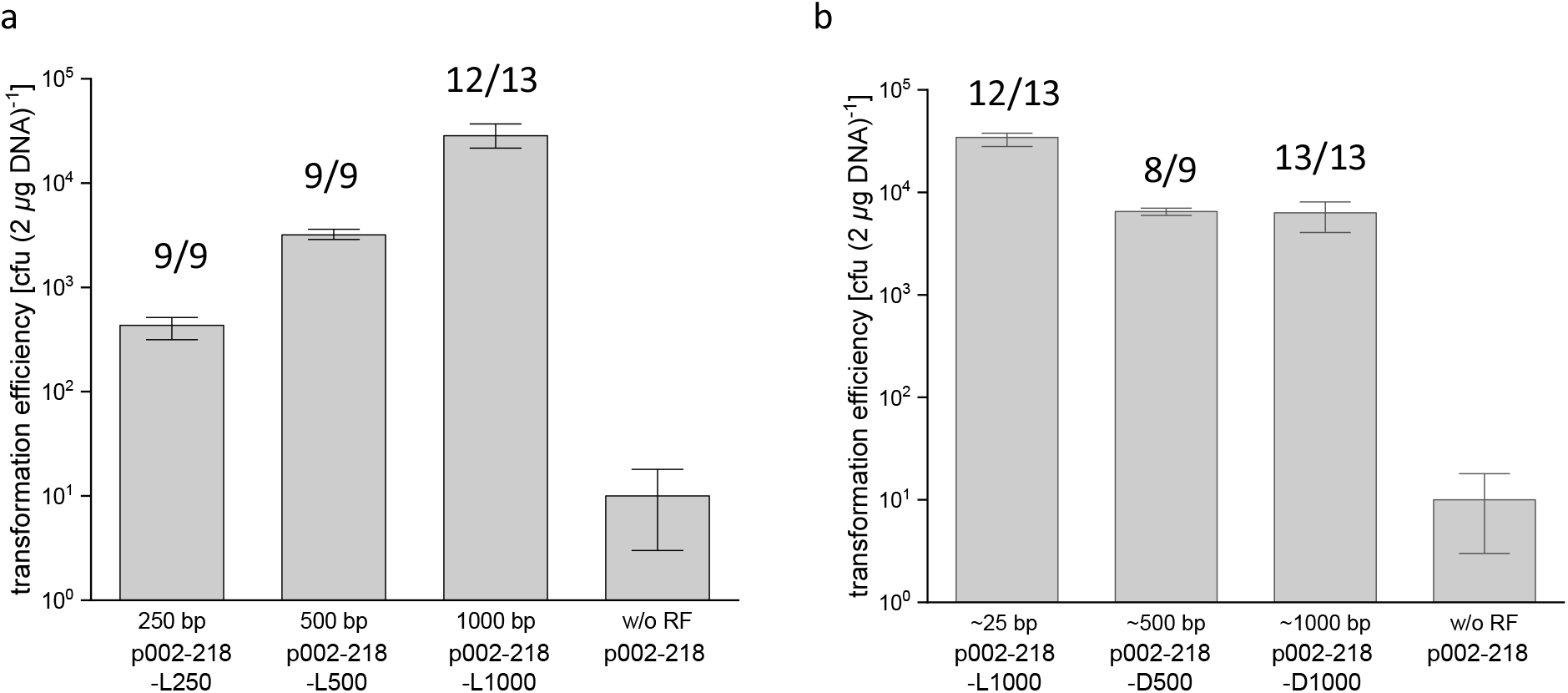
Transformation efficiency and positive rate in relation to the length and position of the RF. a) Variation of the length of the homologous arms flanking the RF, whereby the distance from the RF to DSB is 25 bp. b) Variation of the distance between the RF to the DSB, whereby the length of the homologous arms is 1000 bp. For the 25 bp distance experiment, same plasmid has been used as for the 1000 bp length experiment in a. The error bars for the transformation efficiency represent the standard deviation (n=3). The numbers above the the bars indicate the positive rate (fraction of correctly edited colonies per colonies tested via PCR).

The influence of the distance between the RF and the DSB on the transformation efficiency has been tested by utilizing ~25, ~500, and ~1000 bp distances (Fig. 2b). At ~500 bp distance, the transformation efficiency was 5 times lower than at ~25 bp (two sided *t*-test, *p* < 0.001), but there was no significant difference between ~500 and ~1000 bp distance (two sided *t*-test, *p* > 0.05). Because the *M. maripaludis* JJ strain has an intrinsic *Pst* I restriction/modification system, the host cells can digest the foreign DNA containing unmethylated *Pst* I sites, and one *Pst* I site may cause a 1.6 - 3.4 fold decrease in transformation efficiency^31^. There is one *Pst* I site on the homologous arm on both plasmids (500 bp and 1000 bp), resulting in a lower number of transformants (Fig. 2b). Since the transformation efficiency is the same in the experiments with ~500 bp and with ~1000 bp distances to the DSB, we conclude that the editing efficiency is not significantly affected up to 1000 bp distances.

To test the positive rate, a pair of primers were used to amplify the up- and down-stream of the homologous arms on the chromosome. The PCR products were further digested by the endonuclease NotI. The NotI restriction site was pre-designed between the left and the right RF in order to distinguish the wildtype copies and the edited ones (Fig. S1–S3). The system enabled 89 - 100% positive rate of genome editing (Fig. 2a, Fig. S1–S5). The results indicate that the CRISPR/Cas12a toolbox performs reliable genome editing in *M. maripaludis*. Combining all experiments presented in Fig. 2, 63/66 colonies were correctly edited, which corresponds to an average positive rate of ca. 95%.

### Genome editing via CRISPR/LbCas12a by providing a repair fragment separately

As a variant of the optimized method described above, we have provided the RF as a suicide plasmid which may speed up the work for creating different mutants in parallel. In the section “Quantification of promoter strengths” below, the tested promoters did drive the expression of *uidA* via a *“Promoter-uidA”* expression cassette that has been integrated onto the locus of acetyl-CoA synthetase (MMJJ_09370) via co-transforming the CRISPR/LbCas12 cleavage plasmid with a suicide plasmid containing *“Promoter-uidA”* cassette flanked by 1 kbp up- and down-stream homologous arms. We obtained transformants for this variation with the suicide plasmid, but the transformation efficiency was ca. 10 - 50 fold lower. Although the transformation efficiency is lower in this variant of the standard toolbox, the positive rate appears not to be affected. For all 15 different constructs with suicide plasmid, we randomly tested 3 transformants of each construct, 45/45 of the colonies were found to be positive (Fig. S8)

An additional possibility is to co-transform the CRISPR/LbCas12a cleavage plasmid with a separate RF that was provided as a PCR fragment. Genome editing was successful, but the efficiency was ca. 100-1000 times lower compared to our optimized method with the full p002-g218-L1000 plasmid.

### Replacing a large genome fragment by a heterologous gene via the CRISPR/Cas12a toolbox

To apply the toolbox for the heterologous gene integration, we have replaced the whole flagellum operon (*flaB1B2B3CDEFGHIJ*, length 8.9 kbp) by *uidA* from *E. coli* that codes for a β-glucuronidase. We carried out two different procedures: A) Using one gRNA that generates one lethal DSB on the chromosome, resulting in two long distances from DSB to the repair fragment of 6.4 and 2.5 kbp. B) Cutting on both sides of the *fla* operon by using two different gRNAs, which shortens the distances from DSB to the repair fragment to 0.25 kbp and 3.2 kbp. Both procedures were successful with a similar transformation efficiency (Fig. 3), showing that our toolbox is able to make a large fragment deletion in the genome. The positive rate was lower in procedure A, where one distance was 6.4 kbp (Fig. 3, Fig. S6 and S7), implying that the positive rate is affected when the distance is substantially longer than 1000 bp. A similar observation has been reported for *M. acetivorans:* There, the positive rate dropped significantly when the distance to the DSB was larger than 1000 bp^25^, and the transformation efficiency decreased with increasing distance^25^. Deleting a large fragment by using two gRNAs in order to shorten the distance from RF to DSB demonstrated here may therefore also improve the transformation efficiency and positive rate in other CRISPR systems for methanogenic hosts.

**Figure 3.**
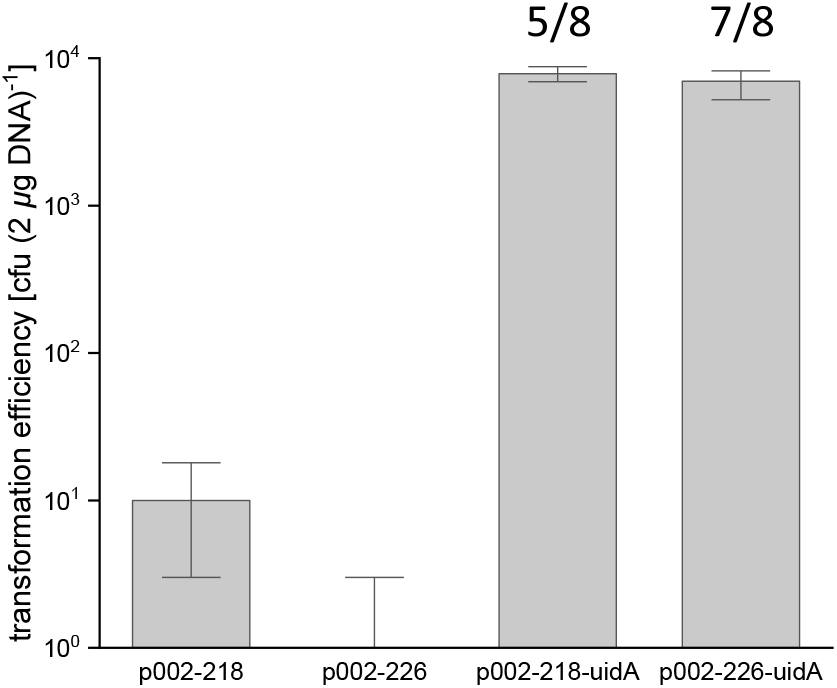
Replacing the whole ORF of the flagellum operon by *uidA* from *E. coli*. p002-218 expresses one gRNA, p002-226 expresses two gRNAs. “uidA” indicates that the RF is present on the corresponding plasmid. The error bars for the transformation efficiency represent the standard deviation (n=3). The numbers above the bars indicate the positive rate (fraction of correctly edited colonies per colonies tested via PCR).

### Genome editing via CRISPR/SpCas9

To expand the versatility and number of usable gRNAs of the CRISPR toolbox, we also constructed a Cas9-based editing plasmid, based on Cas9 from *Streptococcus pyogenes*. To test this additional tool, the gene coding for the alanine dehydrogenase – alanine racemase was successfully replaced by a 4.2 kbp fragment via co-transforming a CRISPR/SpCas9 plasmid with a gRNA and a suicide plasmid containing the 4.2 kbp fragment flanked by 1000 bp up- and down-stream homologous arms. The transformation efficiency was 317 ± 123 cfu (2 *μg* DNA)^-1^ and the positive rate was 8/10 (Fig. S9, Fig. S10). There were two *Pst* I sites on the repair fragment, which can explain the lower transformation efficiency.

### Quantification of promoter strengths

We have tested 15 different promoters: 12 from the closely related methanogen *Methanococcus vannielii* SB, and the three native promoters *P_mcr_, P_mcr_* reverse (*P_mcr_R*) and *P_fla_*. The promotors strengths have been quantified under the two common growth conditions formate or H_2_+CO_2_. All promoters could drive the expression of *uidA* under both growth conditions (formate or H_2_+CO_2_ as substrate), except *P_nif_* and *P_hdrC1_* (Fig. 4). The promotor *P_hdrC1_* allowed gene expression in formate medium, but not in H_2_+CO_2_ medium. The *P_mcr_* promoter is considered as a strong constitutive promoter in methanogens ^32^. Therefore, *P_glnA_*, *P_mtr_*, *P_mcr_*, *P_mcr_JJ_*, *P_fla_JJ_* can be classified as strong promoters in *M. maripaludis*. The *P_nif_* promoter is repressed by the nitrogen regulator, NrpR, and the *P_nif_* becomes highly active once N_2_ is used as a sole nitrogen source or the transcription repressor NrpR is absent^33^. We have deleted *nrpR* in the “*P_nif_-uidA*” strain and found that the strength of *P_nif_* increased significantly (2670 ± 58 nmol min^-1^ OD_600_^-1^) (two sided *t*-test, *P* < 0.001) when NrpR was absent in *M. maripaludis*. Thus, the “*ΔnrpR-P_nif_*” expression may be used for target genes requiring a very strong expression.

**Figure 4.**
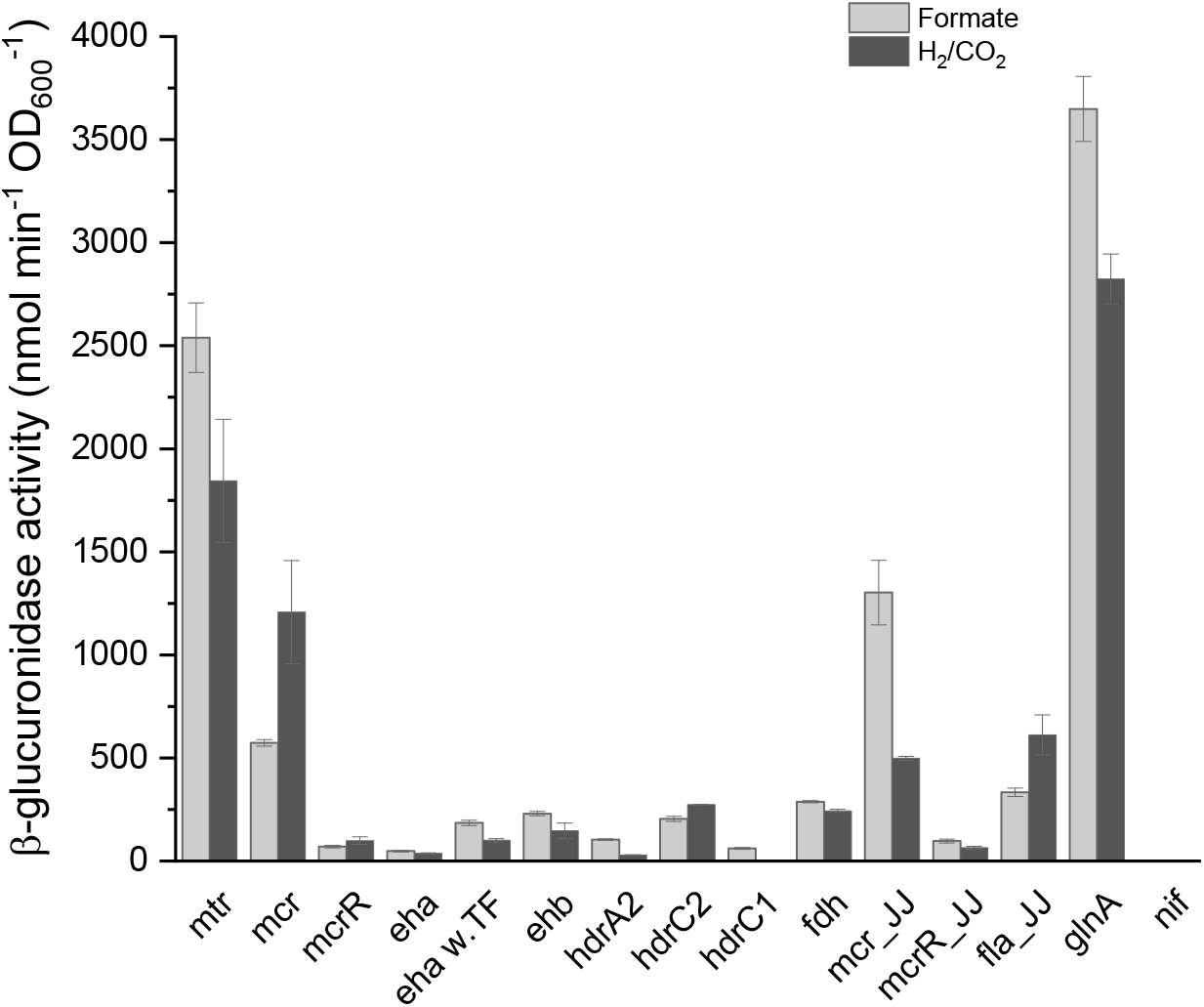
Quantification of promoter strengths for the two different growth conditions formate or H_2_+CO_2_, measured after the culture has reached OD_600_ = ca. 0.5. The promoters mcr_JJ, mcrR_JJ and fla_JJ are from *M. maripaludis* JJ. The remaining promoters are from *Methanococcus vannielii* SB. Error bars represent the standard deviation (n=3).

The majority of the tested promoters had a similar strength on both growth-conditions, formate and H_2_+CO_2_. The *P_eha_* promoter that drives the expression of the energy conservating hydrogenase A, *eha*, in *M. vannielii* was weak in *M. maripaludis*. Notably, the *P_eha_* did not directly express the subunits of *eha*. The first gene driven by *P_eha_* is a putative transcription regulator. Thus, we also tried to express *uidA* after the the putative transcription regulator (*P_eha w.TF_*). The expression level of UidA was significantly higher with *P_eha w.TF_* than *P_eha_* (two sided *t*-test, *p* < 0.01) in both growth conditions. These results imply that the transcription regulator might be involved in the regulation of *eha*.

The *P_glnA_* promoter from *M. vannielii* was unexpectedly strong in *M. maripaludis* despite of the presence of NrpR and ammonium in the medium. In order to exclude a potential mutation, we have sequenced the *P_glnA_-uidA* expression cassette from the genome of the engineered *M. maripaludis* strain, but no mutation could be found. We have also repeated the transformation and re-picked several different transformants to test the UidA activity. All transformants showed a high expression level of UidA. Thus, we can conclude that *P_glnA_* from *M. vannielii* is a strong promoter in *M. maripaludis*. The native *P_glnA_* promoter has a basal constitutive expression level in *M. maripaludis* when ammonium is present^34^. Although the *glnA* operator from *P_glnA_* promoter is the same as the one from *M. maripaludis*, the expression level of “*P_glnA_ -uidA*” appears to be the highest among all promoters tested.

## CONCLUSION

*M. maripaludis* already has genetic tools and efficient transformation protocols^17,31,28^ for standard applications, but many colonies may need to be screened to obtain the desired genotype by the classic pop-in-pop-out genome editing technique^17^, especially when the targets to be engineered affect cell growth^32^. As a solution, we have established a reliable CRISPR/Cas12a toolbox in *M. maripaludis* for gene knock-in and knock-out with positive rate of ca. 95%. Our system only requires one round of homologous recombination, and no merodiploid is formed, which lowers the workload of genome editing and increases the success rate. The option of providing the repair fragment separately as a suicide plasmid or as a PCR template may further speed up the editing progress. Since Cas12a has intrinsic ribonuclease activity that can process multiplex gRNA in one transcript without extra ribonuclease^26,27^, it can be convenient to express two gRNAs via the CRISPR/LbCas12a system, saving time and cost of plasmid construction.

Proteins can be expressed heterologously in *M. maripaludis* via stable integration of the corresponding genes. Utilizing *M. maripaludis* as an expression host may be advantageous for proteins that are difficult to be expressed in *E. coli*, such as formate dehydrogenases^16^, methyl-coenzyme M reductases^35^, or heterodisulfide reductases^36^.

Although different promoters have already been studied and used for synthetic biology in *M. maripaludis*^37,14,38,33^, there has not been a uniform system to compare their strengths, which is now available and allows to balance metabolic fluxes in engineered *M. maripaludis* strains.

## MATERIALS AND METHODS

### Strain and plasmids

All plasmids and strains used in this study are listed in Table S1 and Table S2. The links of plasmid maps are listed in Table S3. *M. maripaludis* JJΔupt^28^ and plasmid pLW40^29^ are gifts from Prof. Kyle Costa, University of Minnesota. Plasmid pMEV4^4^ was kindly provided by Prof. William B Whitman, University of Georgia. *M. maripaludis* S2^39^ was kindly provided by Prof. John Leigh and Dr. Thomas Lie, University of Washington. NEB5α (New England Biolabs) was used for plasmid construction. The plasmid pMM002P and pMM005 was constructed by Gibson assembly^40^. The construction procedure and primers of pMM002P and pMM005 are described in Table S4 and S5. All the cleavage plasmids were constructed as follows: (1) for LbCas12a gRNA, the forward primer was designed as “5’-AGAT+(23nt guide sequence)”, the reverse primer was designed as “5’-TATC+(23nt reverse complement guide sequence)”; for SpCas9 gRNA, the forward primer was designed as “5’-AGTG+(20nt guide sequence)”, the reverse primer was designed as “5’-AAAC+(20nt reverse complement guide sequence) (2) the two primers which contain the gRNA and have 4 nt overhang at 5’-end were annealed (3) the annealing product was ligated with *PaqC* I (New England Biolabs)-digested pMM002P or pMM005. The guide sequences was designed on CHOPCHOP (https://chopchop.cbu.uib.no/)^41^. The RF was inserted into the corresponding cleavage plasmid at the *Mre* I restriction site. The rest primers used in this study are listed in Table S6.

### Medium and cultivation conditions

Luria–Bertani medium with 50 mg L^-1^ ampicillin was used for plasmid construction and amplification. McC medium was used for *M. maripaludis* strain cultivation with 2.8 bar 80% H_2_/20% CO_2_ in the headspace^28^. The culture tubes were incubated at 37°C at 200 rpm. McFC medium was used when formate was used as a carbon source^42^ The culture tubes were incubated at 37°C without shaking. 2.5 *μ*g mL^-1^ puromycin or 0.25 mg mL^-1^ 6-azauracil was added where required.

### *M. maripaludis* transformation

The natural transformation protocol was used as previously described^28^. Briefly, 5 mL of *M. maripaludis* culture was grown overnight to an OD_600_ between 0.7 and 1.2. 2 *μ*g of DNA was added into the culture directly, followed by flushing the headspace with 80% H_2_/20% CO_2_; for 0.5 min and pressurizing the cultivation to 2.8 bar. The tube was then incubated in a shaker at 37°C at 200 rpm for 4 h and the cells plated on McC medium.

### Removal of the CRISPR/Cas plasmid

The *M. maripaludis* strains containing the CRISPR/Cas toolbox plasmid was grown in 5 mL of McC medium without antibiotics to an OD_600_ between 0.7 and 1.2. Then 100 *μ*L of the preculture was inoculated in another 5 mL of McC medium without antibiotics for overnight. On the next day, one drop of culture was streaked on the McC medium with 6-azauracil plate. After 3-5 days, several individual colonies were picked and streaked on another McC medium 6-azauracil plate for purification.

### β-glucuronidase activity measurements

4-Nitrophenyl β-*D*-glucuronide (4-NPG, CAS no. 10344-94-2) (Sigma-aldrich) was prepared in 50 mM sodium phosphate buffer (Na-PB) (pH=7.0) as a stock solution (10 mg mL^-1^). The detailed procedure was as follows: the cells were grown to an OD_600_ of ca 0.5; the exact OD_600_ was measured (BioPhotometer plus, Eppendorf); 1 mL of cell culture was pelleted via centrifugation; the cell pellet was resuspended in 500 *μ*L of Na-PB; around 30 μl glass beads were added; the tube was vortexed for 5 min to break the cells thoroughly; the tube was centrifuged and the supernatant collected; the sample was diluted to 500 *μ*L in Na-PB; the sample incubated at 37°C for 20 minutes; 40 *μ*L 4-NPG was added; reaction mixture was kept at 37°C for 15 minutes to react; 400 *μ*L 200 mM Na_2_CO_3_ was added to stop the reaction; the absorbance at 405 nm was read. The specific activities have been calculated using β-glucuronidase from *E. coli* K12 (Product no. 3707580001, Sigma-Aldrich) as the standard, using the conversion factor of 398 nmol min^-1^. The standard curve is shown in Fig. S11.

## AUTHOR CONTRIBUTIONS

JB and SS conceived the study and wrote the manuscript. JB and EDM performed the experiments. All authors commented and approved the final version of the manuscript. All authors declare no conflict of interest.

## ACKNOWLEDGEMENTS

We thank the summer-students Han Le, An Nguyen and Pradhuman Jetha for contributing to the plasmid construction work and *M. maripaludis* transformation. We thank Prof. Kyle Costa for providing strain JJΔupt and plasmid pLW40, Prof. William B. Whitman for providing plasmid pMEV4,pMEV4mTs, and strain *M. voltae* A3, and Prof. John Leigh and Dr. Thomas Lie for providing strain S2. Prof. Michael Rother, Prof. Qunxin She and Prof. Dipti Nayak are acknowledged for helpful discussions. We thank Dr. Norman Adlung, Dr. Ingemar von Ossowski and Dr. Vera Jäger for the critical comments on the manuscript. We thank the Novo Nordisk Foundation for funding (grant NNF19OC0054329 to S. S.).

---

**Table S1.**
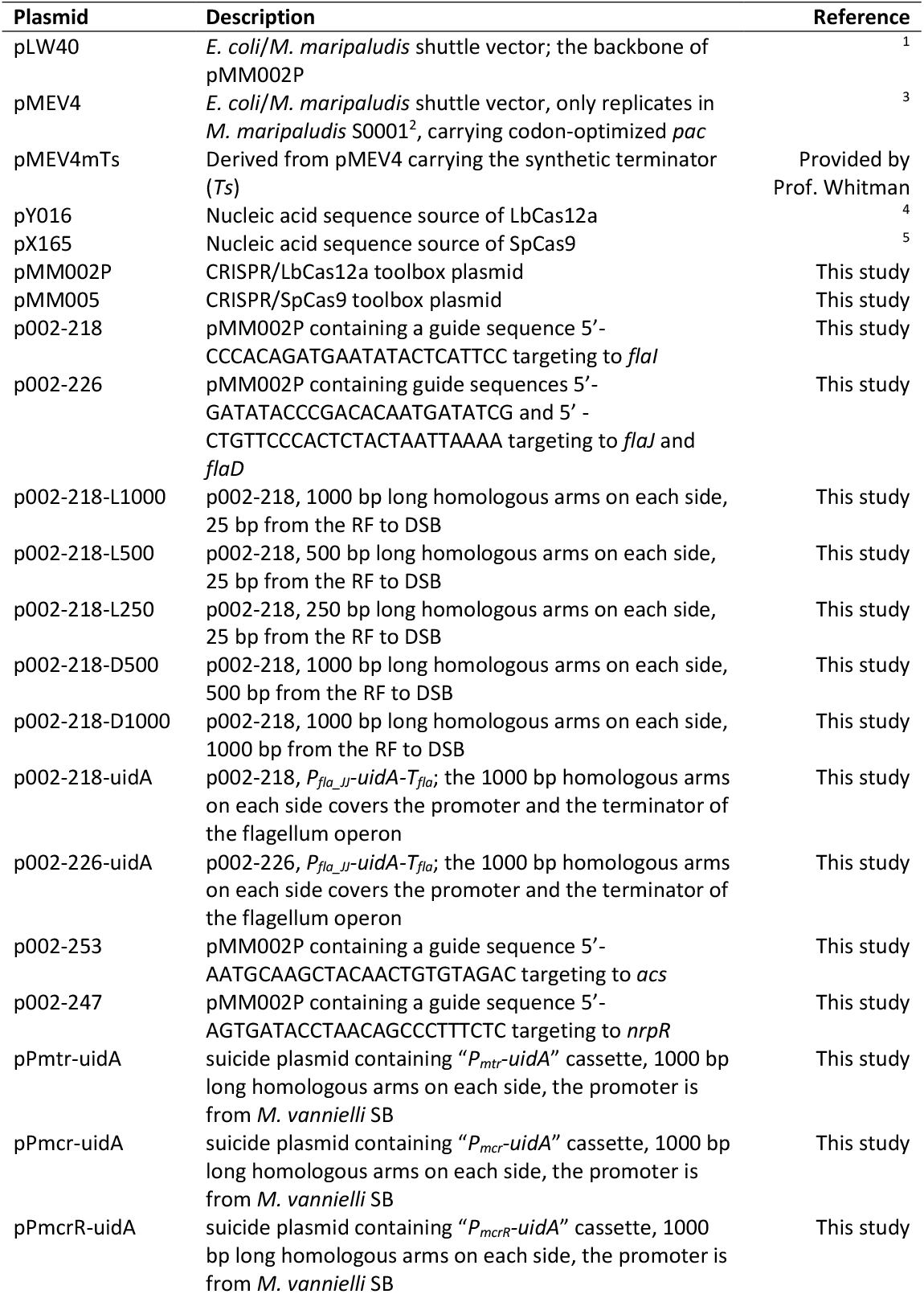
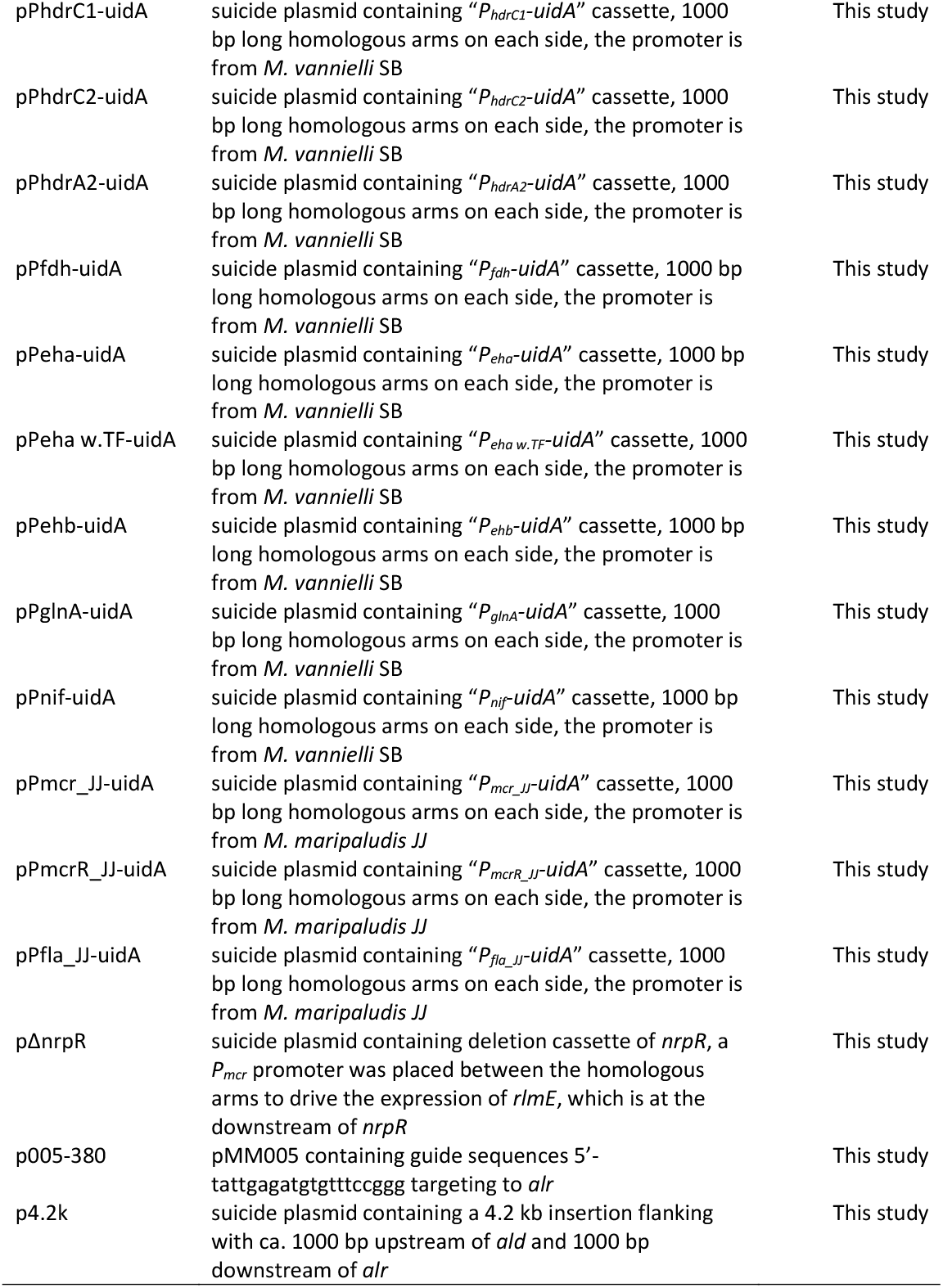
Plasmid list

**Table S2.**
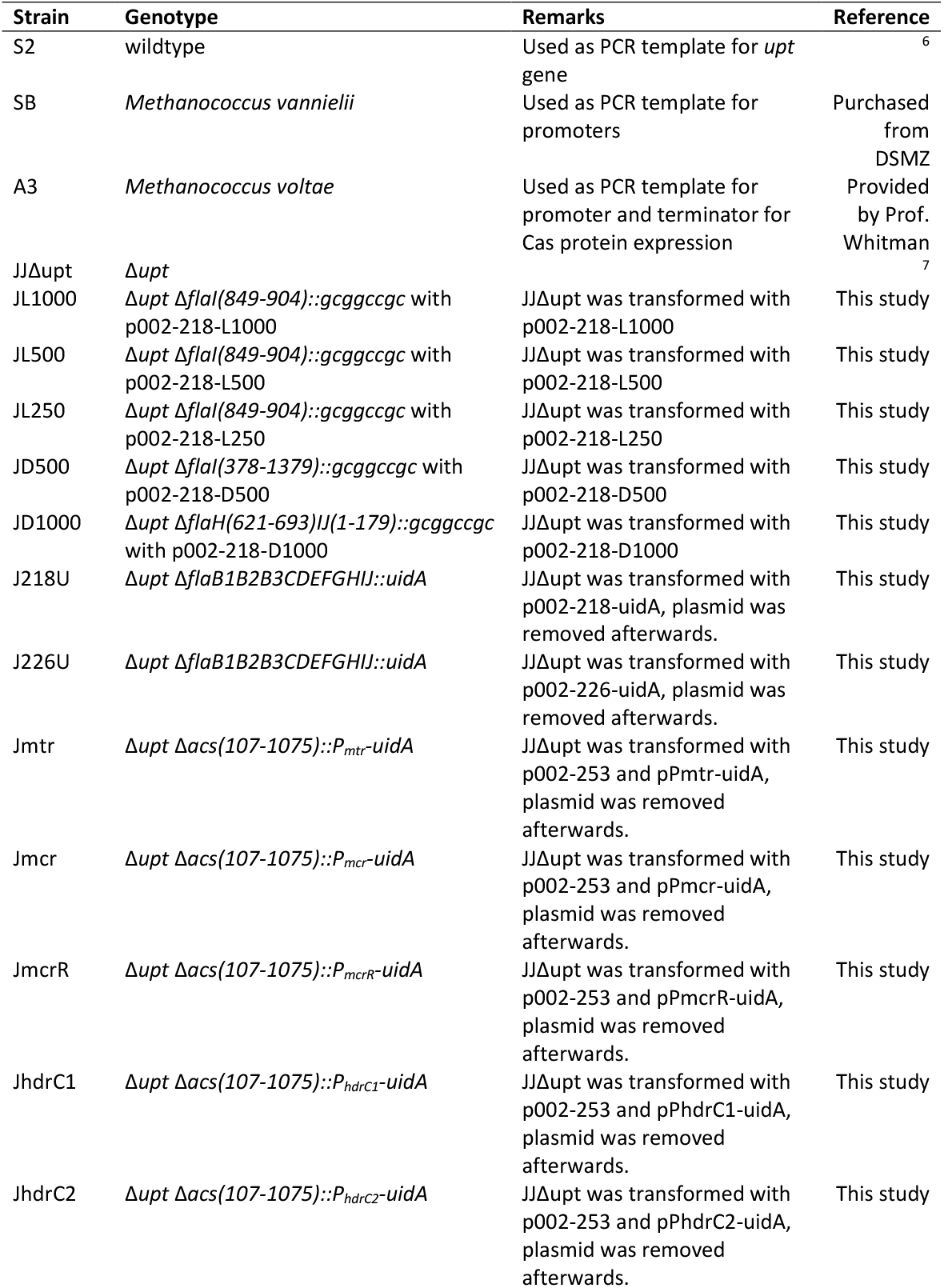
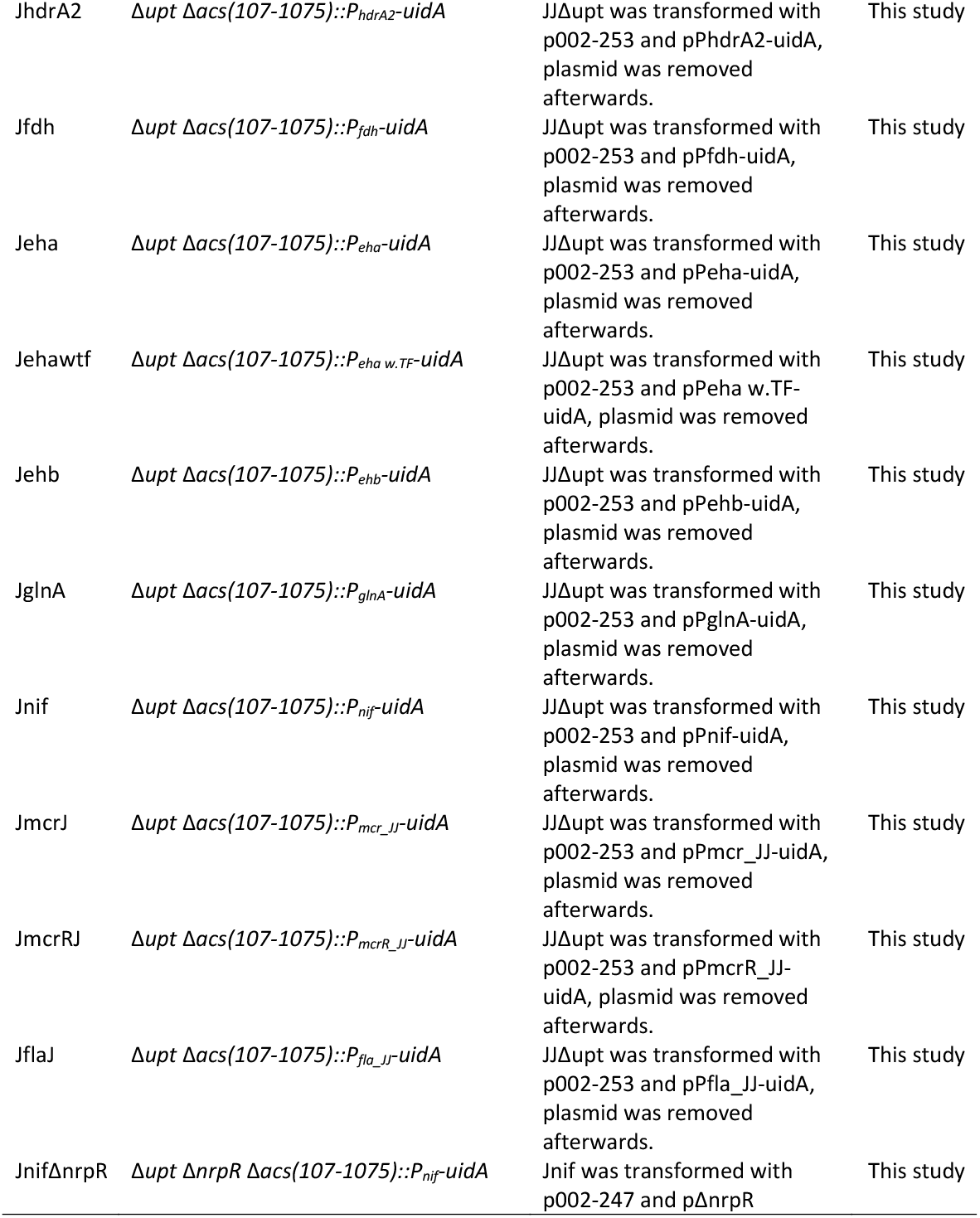
Strain list

**Table S3.**
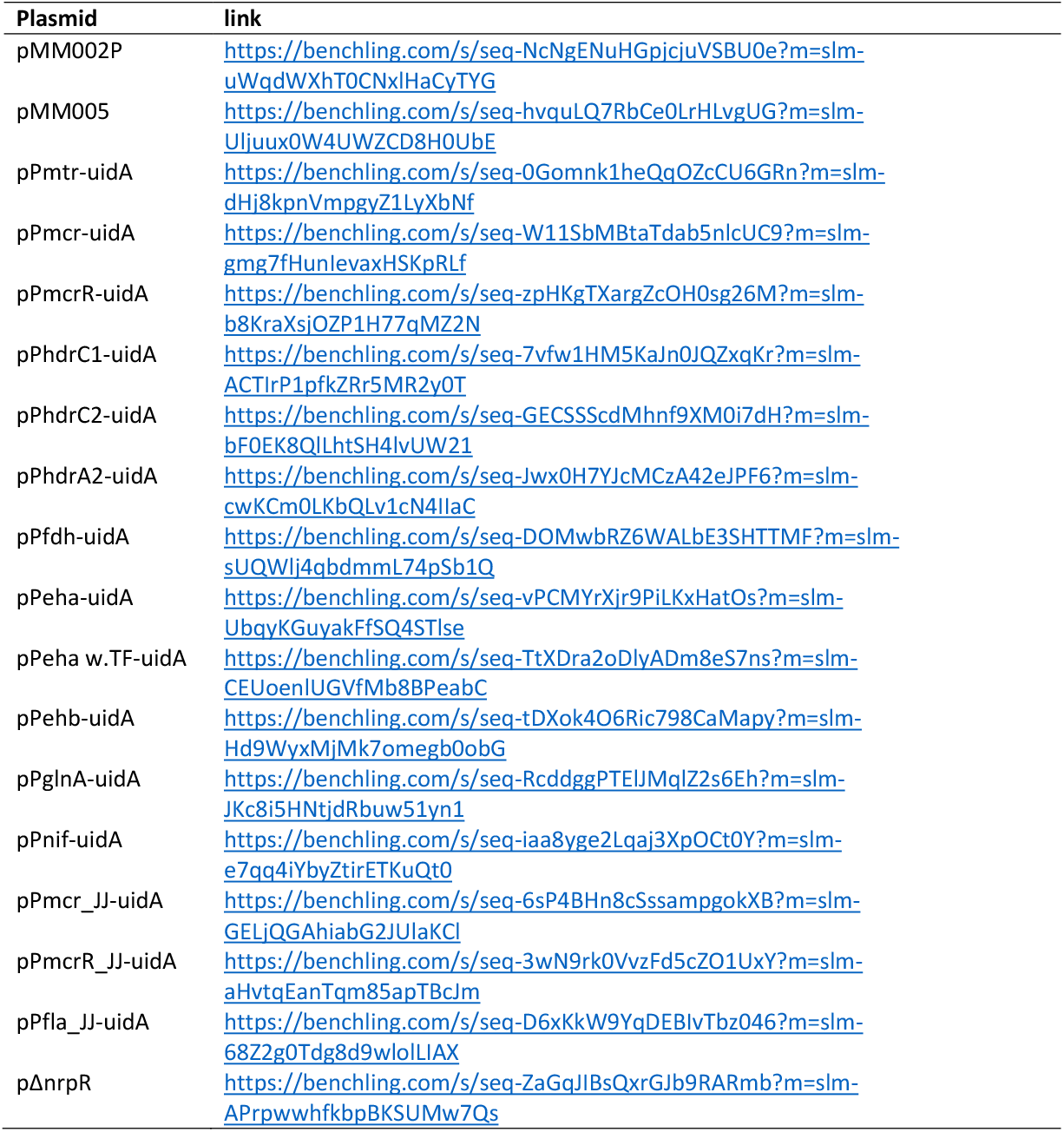
plasmid maps

**Table S4.**
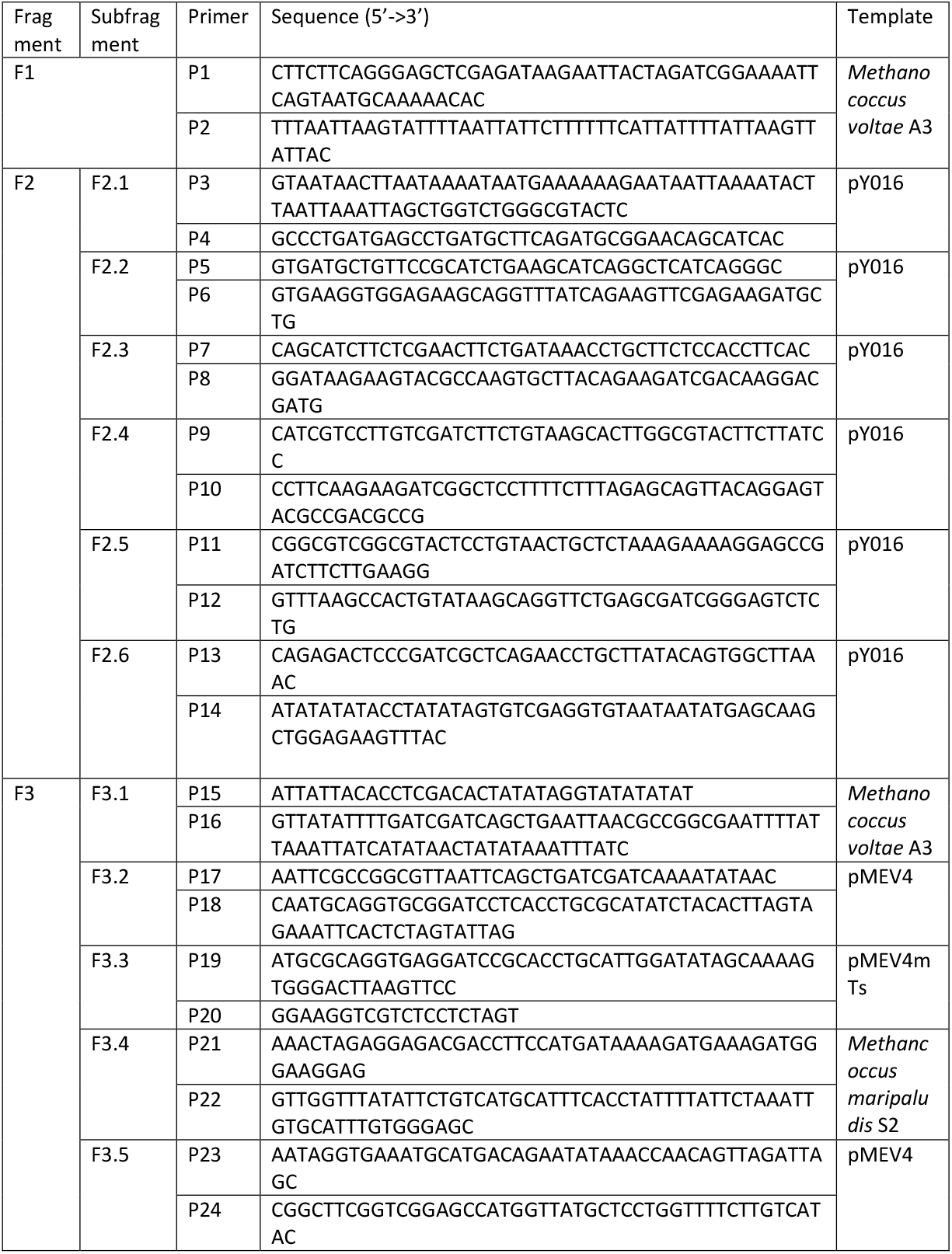
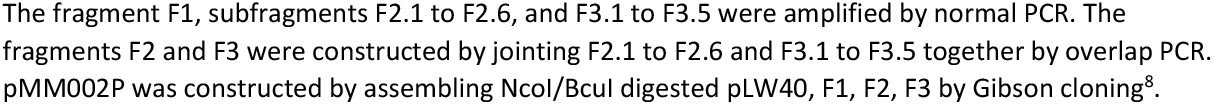
Primers used for pMM002P construction

**Table S5.**
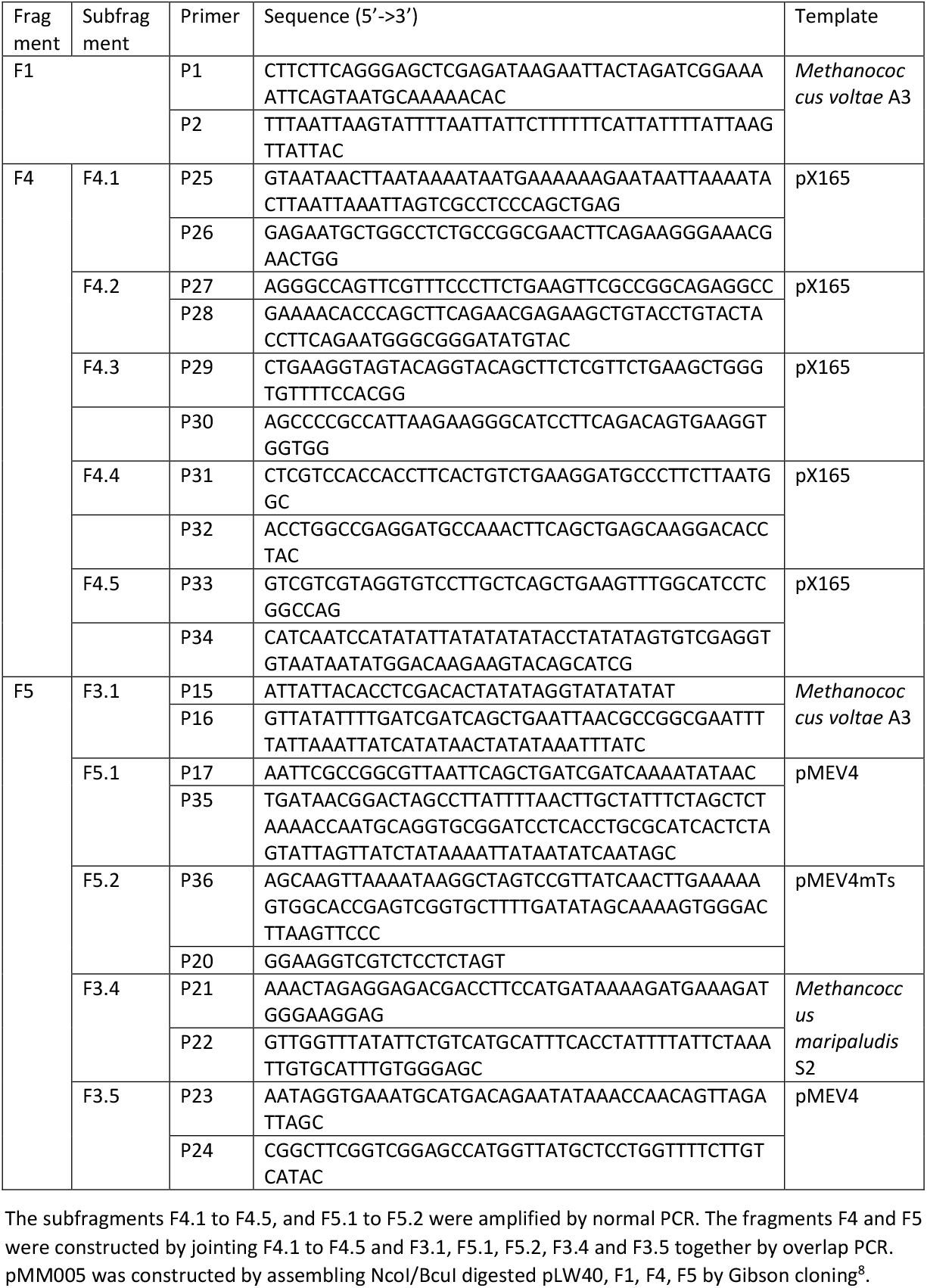
Primers used for pMM005 construction

**Table S6.**
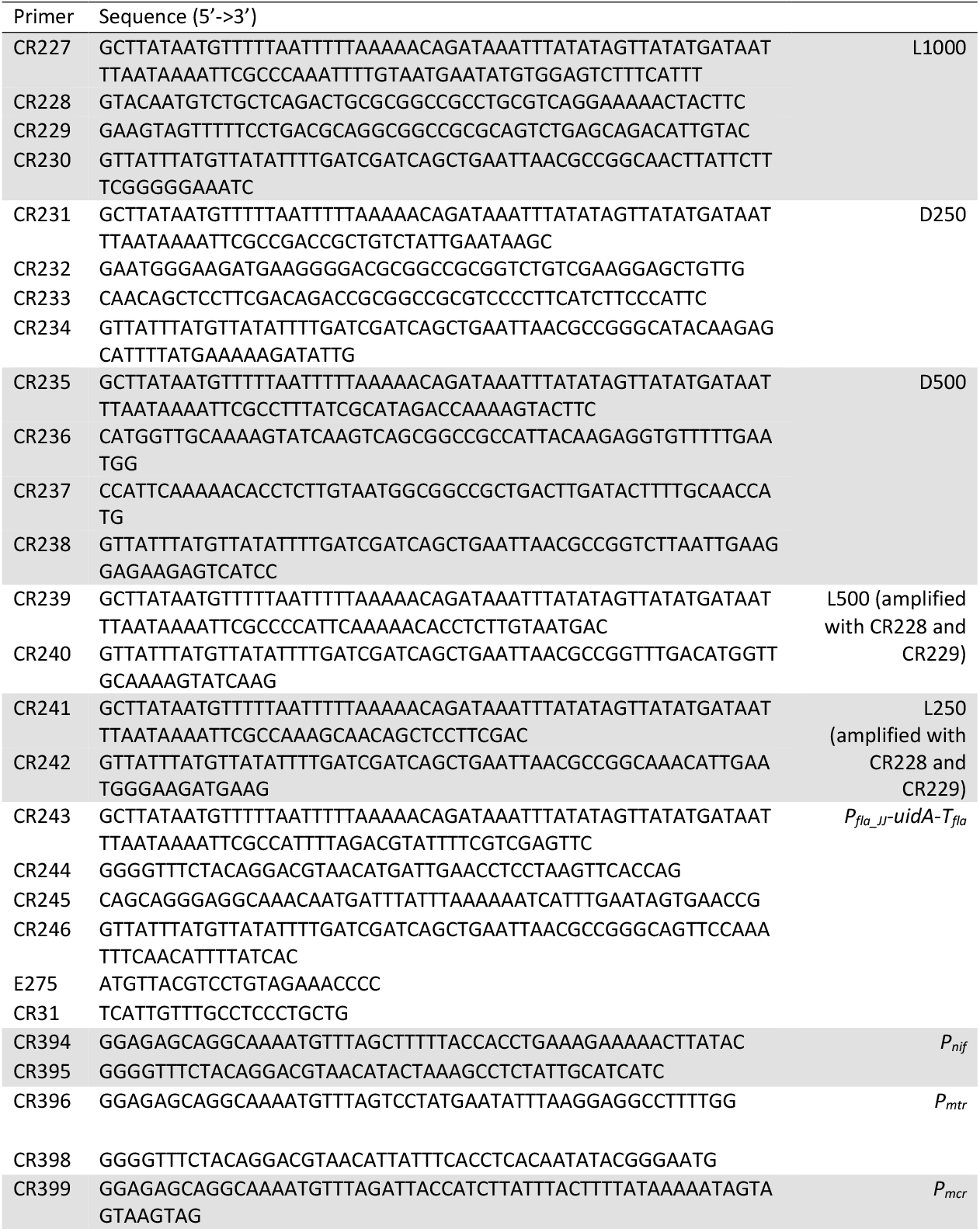
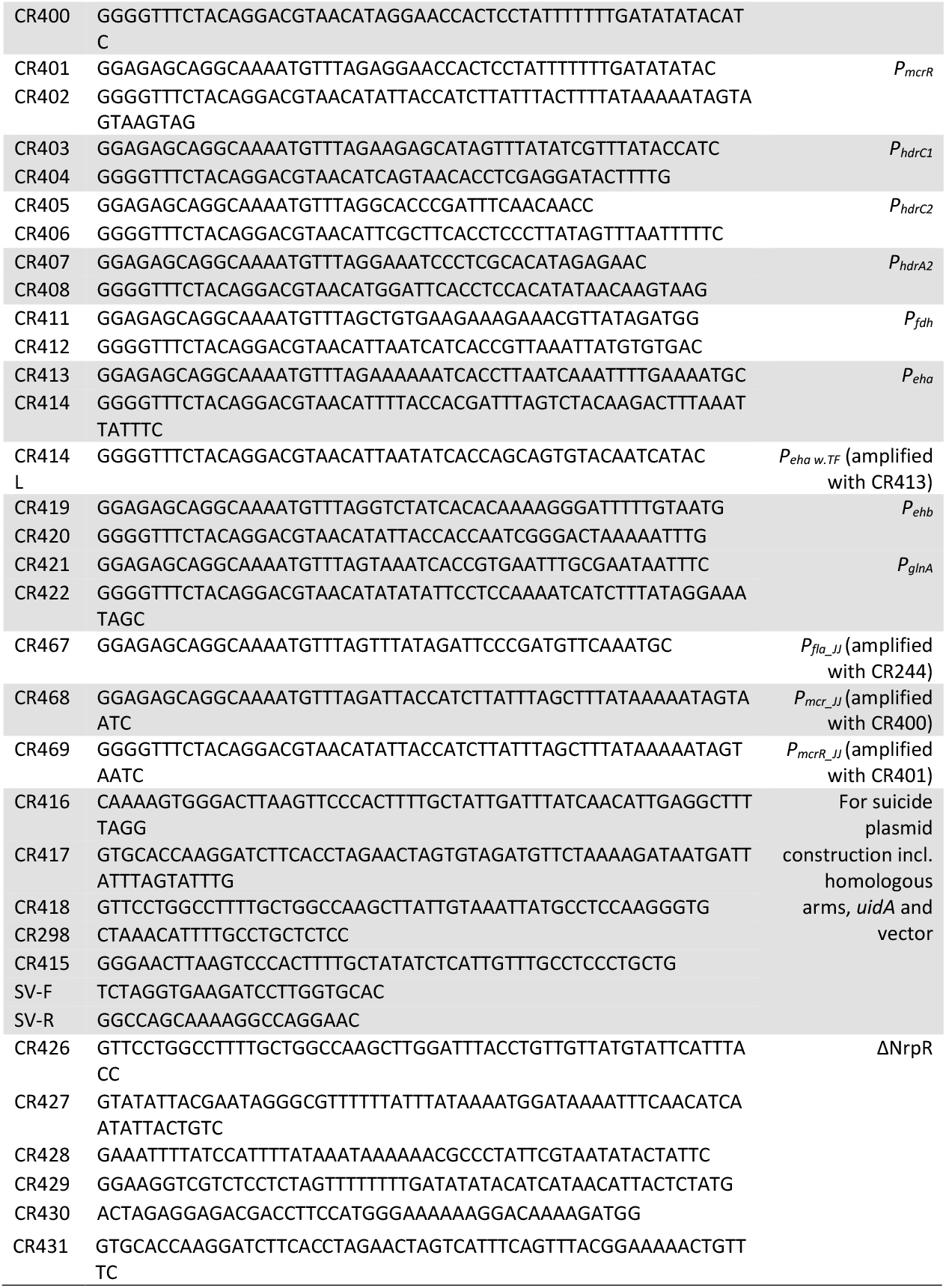
The rest of primers used in this study

**Figure S1.**
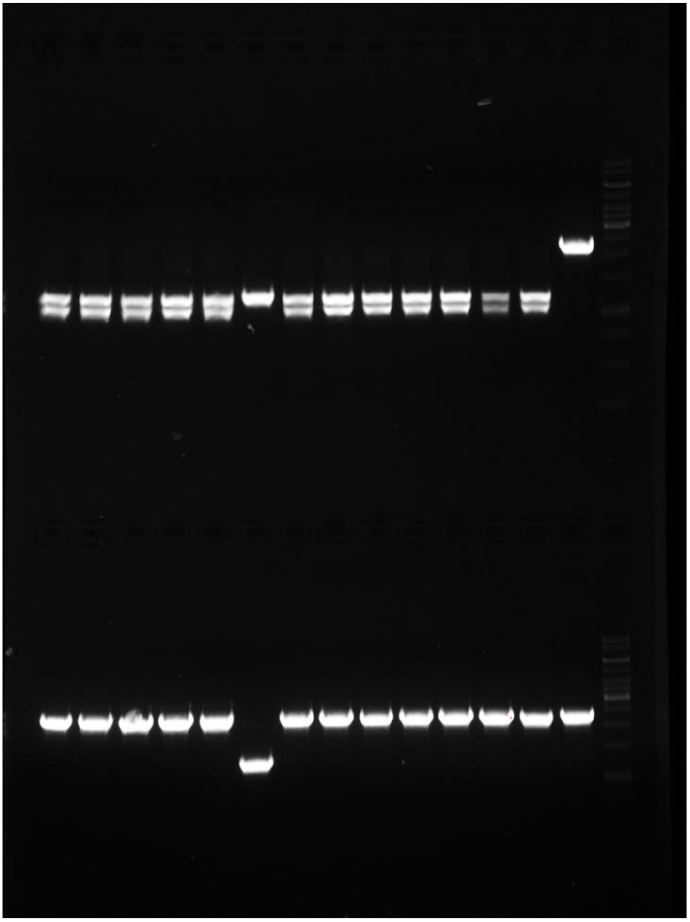
Colony PCR results for genome editing with the plasmid p002-218-L1000. Upper row: PCR products digested by NotI. Lower row: Undigested PCR products. Lanes 1-13: individual colonies. Lane 14: wildtype. Lane 15: DNA ladder (Thermo Scientific GeneRuler 1 kb DNA ladder #SM0311). The primers amplify from 225 bp upstream and 69 bp downstream of the RF on the genome. The expected size of the PCR products of the edited transformants is 2302 bp, that of the wildtype is 2350 bp. The NotI digested PCR products (upper row) are expected to have two bands: 1227 bp and 1075 bp, if editing took place. The PCR product of the wildtype genome is not digestible by NotI.

**Figure S2.**
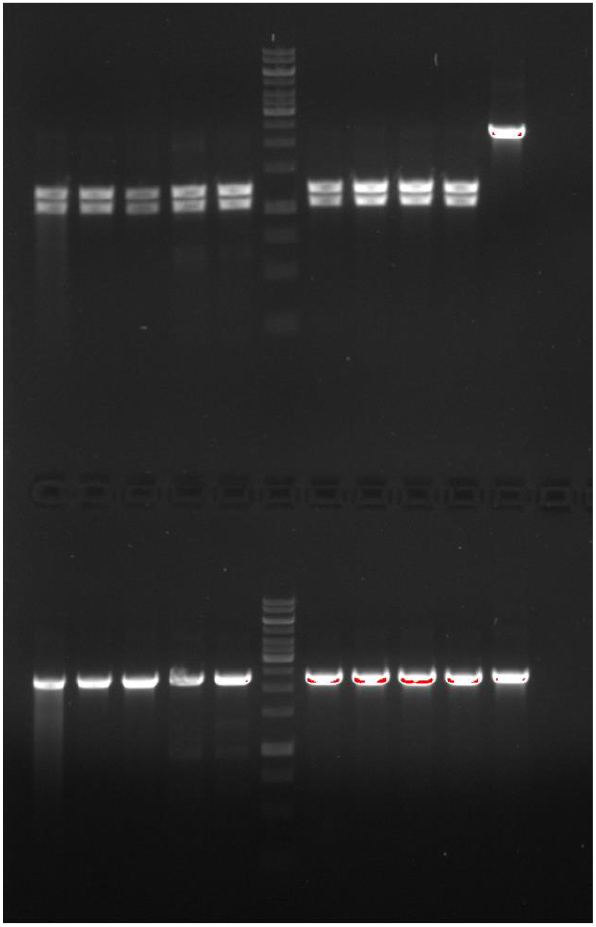
Colony PCR result of p002-218-L500 edited colonies. Upper row: PCR products digested by NotI. Lower row: Undigested PCR products. Lanes 1-5, 7-10: individual colonies. Lane 11: wildtype. Lane 6: DNA ladder (Thermoscientific GeneRuler 1 kb DNA ladder #SM0311). The primers amplify from 725 bp upstream and 569 bp downstream of the RF on the genome. The expected size of the PCR products of the edited transformants is 2302 bp, that of the wildtype is 2350 bp. The NotI digested PCR products (upper row) are expected to have two bands: 1227 bp and 1075 bp, if editing took place. The PCR product of the wildtype genome is not digestible by NotI.

**Figure S3.**
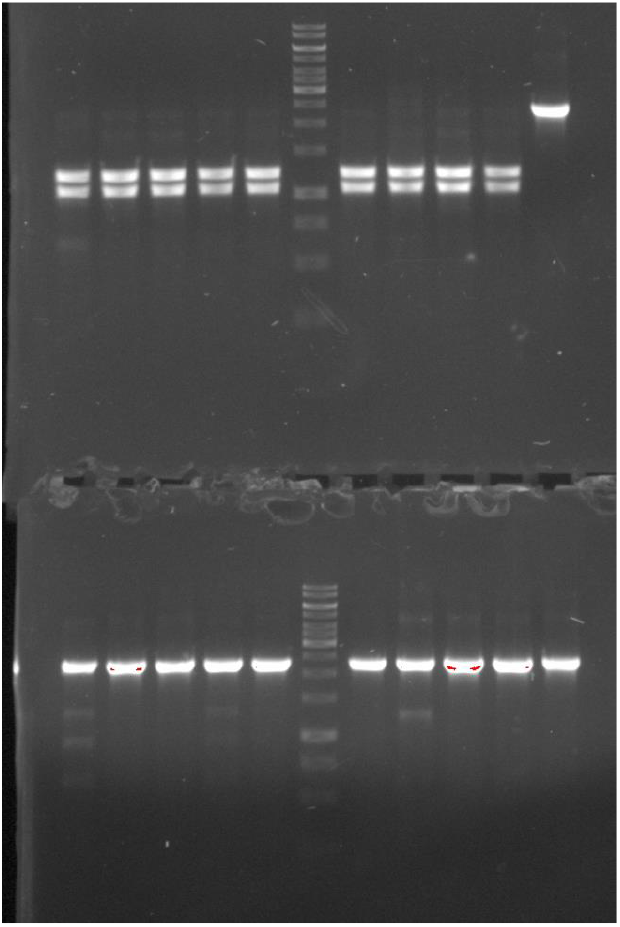
Colony PCR result of p002-218-L250 edited colonies. Upper row: PCR products digested by NotI. Lower row: Undigested PCR products. Lanes 1-5, 7-10: individual colonies. Lane 11: wildtype. Lane 6: DNA ladder (Thermoscientific GeneRuler 1 kb DNA ladder #SM0311). The primers amplify from 725 bp upstream and 569 bp downstream of the RF on the genome. The expected size of the PCR products of the edited transformants is 2302 bp, that of the wildtype is 2350 bp. The NotI digested PCR products (upper row) are expected to have two bands: 1227 bp and 1075 bp, if editing took place. The PCR product of the wildtype genome is not digestible by NotI.

**Figure S4.**
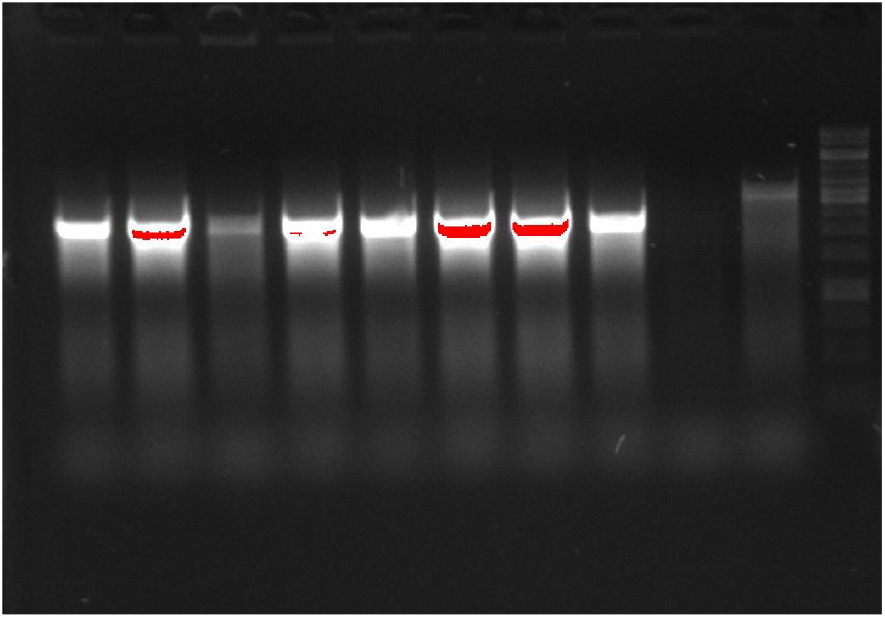
Colony PCR result of p002-218-D500 edited colonies. Lane 1-9: individual colonies. Lane 10: wildtype. Lane 11: DNA ladder (Thermoscientific GeneRuler 1 kb DNA ladder #SM0311). The primers amplify from 215 bp upstream and 57 bp downstream of the RF on the genome. The expected size of the PCR products of the edited transformants is 2280 bp, that of the wildtype is 3274 bp.

**Figure S5.**
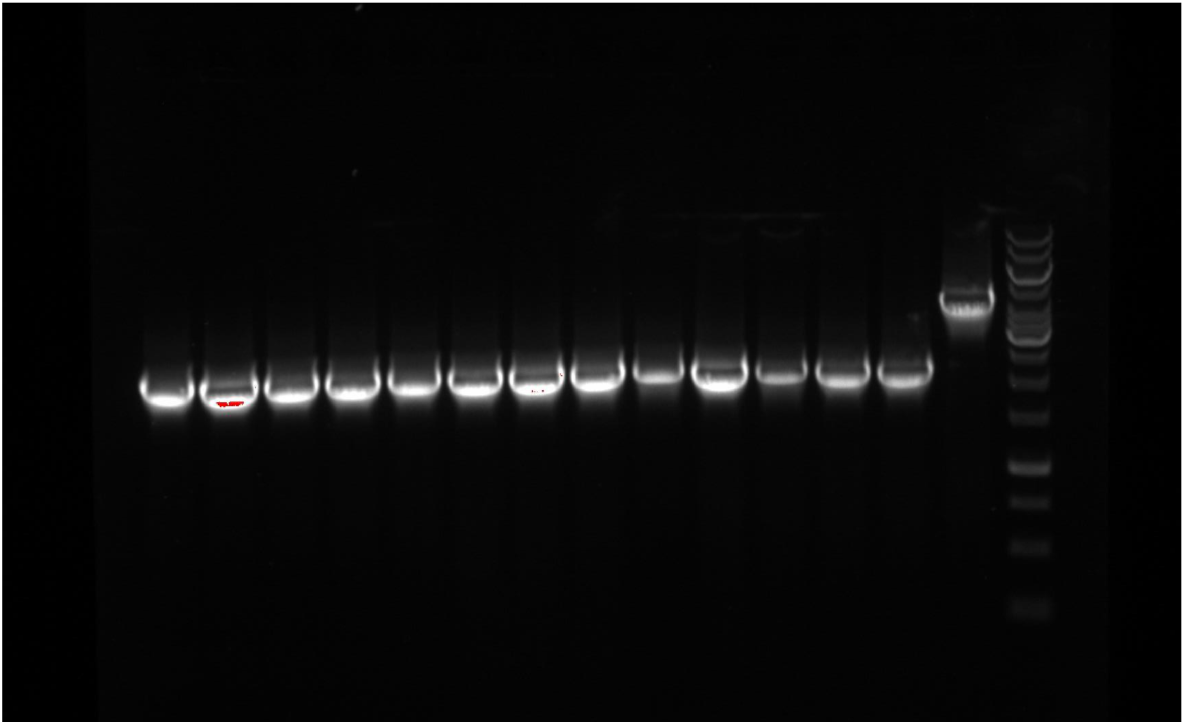
Colony PCR result of p002-218-D1000 edited colonies. Lane 1-13: individual colonies. Lane 14: wildtype. Lane 15: DNA ladder (Thermoscientific GeneRuler 1 kb DNA ladder #SM0311). The primers amplify from 64 bp upstream and 102 bp downstream of the RF on the genome. The expected size of the PCR products of the edited transformants is 2174 bp, that of the wildtype is 4078 bp.

**Figure S6.**
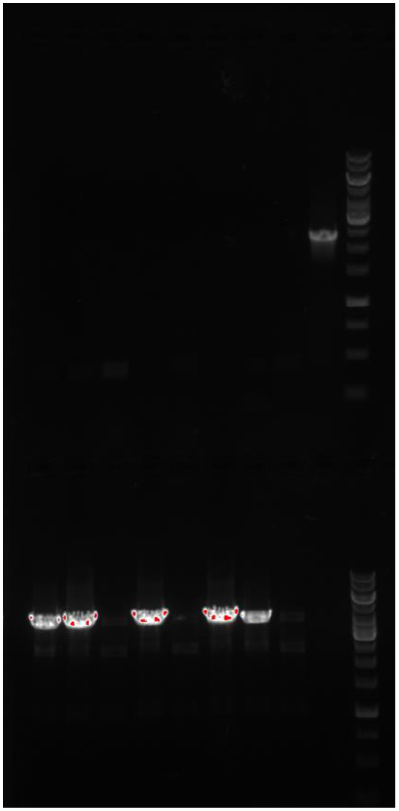
Colony PCR result of p002-218-uidA edited colonies. Lane 1-8: individual colonies. Lane 9: wildtype. Lane 10: DNA ladder (Thermoscientific GeneRuler 1 kb DNA ladder #SM0311). Two pairs of primers were used to check whether the whole flagellum ORF were replaced with *uidA*. The upper row is the amplicon of the part of the flagellum ORF. The expected size of the amplicon of the edited transformants and wildtype genome is no band and 2350 bp, respectively. The lower row is the amplicon from 64 bp upstream and 86 bp downstream of the RF on the genome. The expected size of the PCR products of the edited transformants and wildtype genome is 3905 bp and 10983 bp, respectively. The 10983 bp band for the wildtype genome is not visible because our polymerases were not able to amplify a fragment larger than 8 kb.

**Figure S7.**
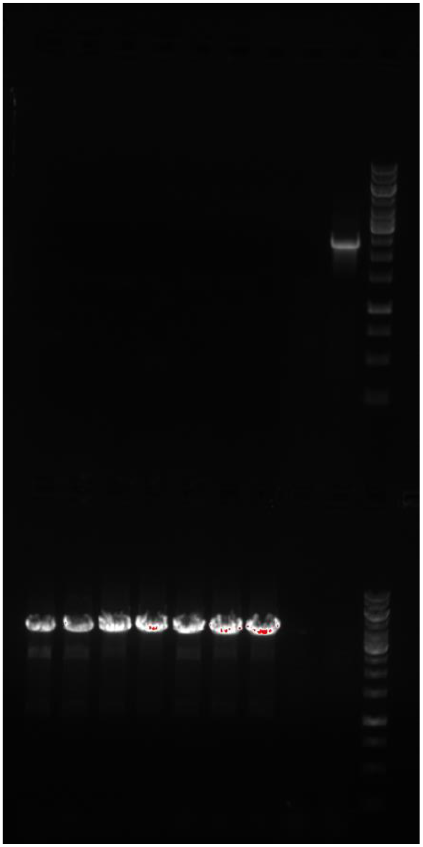
Colony PCR result of p002-226-uidA edited colonies. Lane 1-8: individual colonies. Lane 9: wildtype. Lane 10: DNA ladder (Thermoscientific GeneRuler 1 kb DNA ladder #SM0311). Two pairs of primers were used to check whether the whole flagellum ORF were replaced with *uidA*. The upper row is the amplicon of the part of the flagellum ORF. The expected size of the amplicon of the edited transformants and wildtype genome is no band and 2350 bp, respectively. The lower row is the amplicon from 64 bp upstream and 86 bp downstream of the RF on the genome. The expected size of the PCR products of the edited transformants and wildtype genome is 3905 bp and 10983 bp, respectively. The 10983 bp band for the wildtype genome is not visible because our polymerases were not able to amplify a fragment larger than 8 kb.

**Figure S8.**
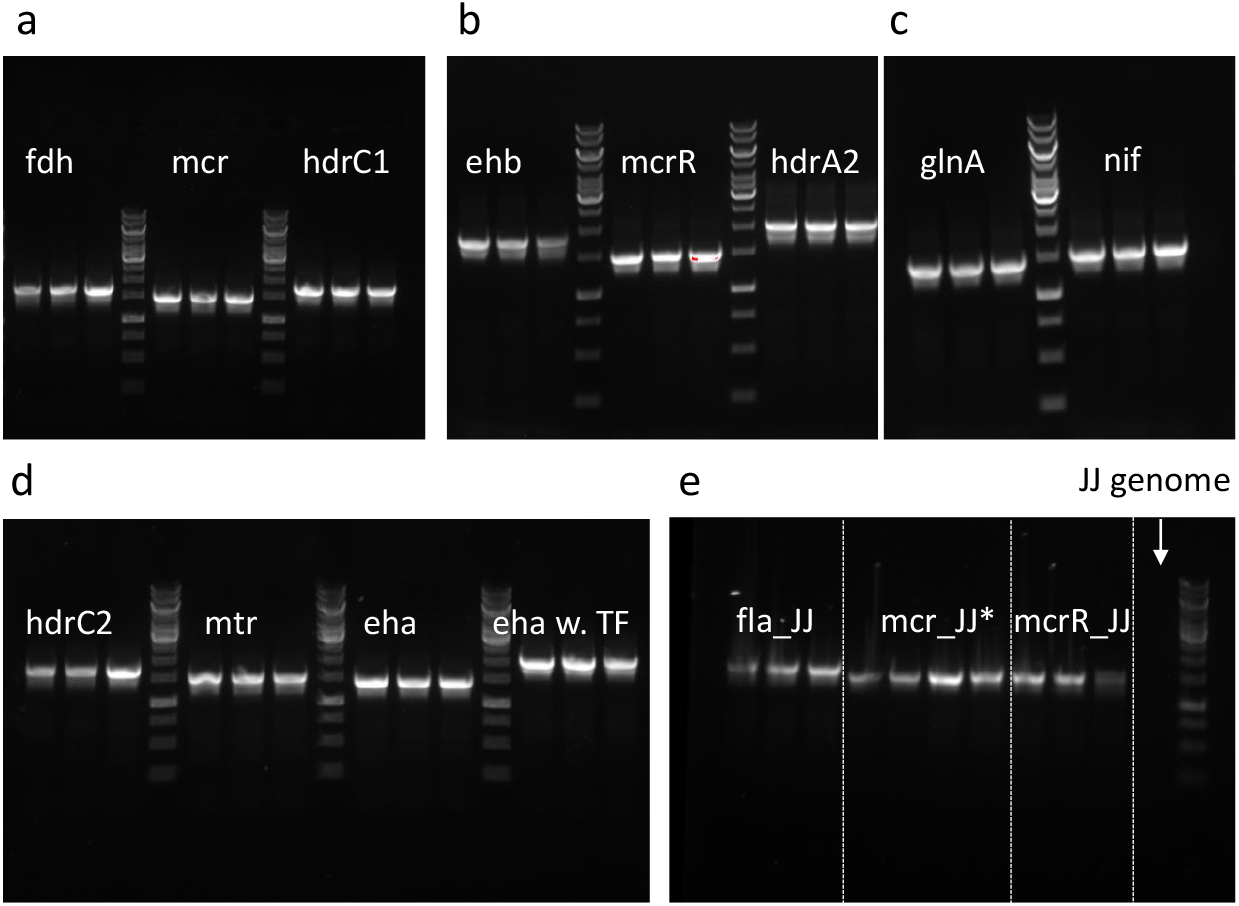
Colony PCR result of co-transforming a CRISPR/LbCas12a plasmid and a suicide plasmid with *“Promoter-uidA”* cassette flanking with 1000 bp homologous repair fragment. One pair of primers was used to check whether the *“Promoter-uidA”* cassette was successfully inserted onto the genome. The forward primer amplifies from 56 bp upstream of the up homologous arm and reverse primer amplifies from the internal of the *uidA* gene. The expected size of the amplicon of the edited transformants is 1189 bp + the size of the promoter. No band should be obtained from the wildtype genome. *In Fig. S8e, the first mcr_JJ sample was accidentally loaded on both lanes: 4 and 5.

**Figure S9.**
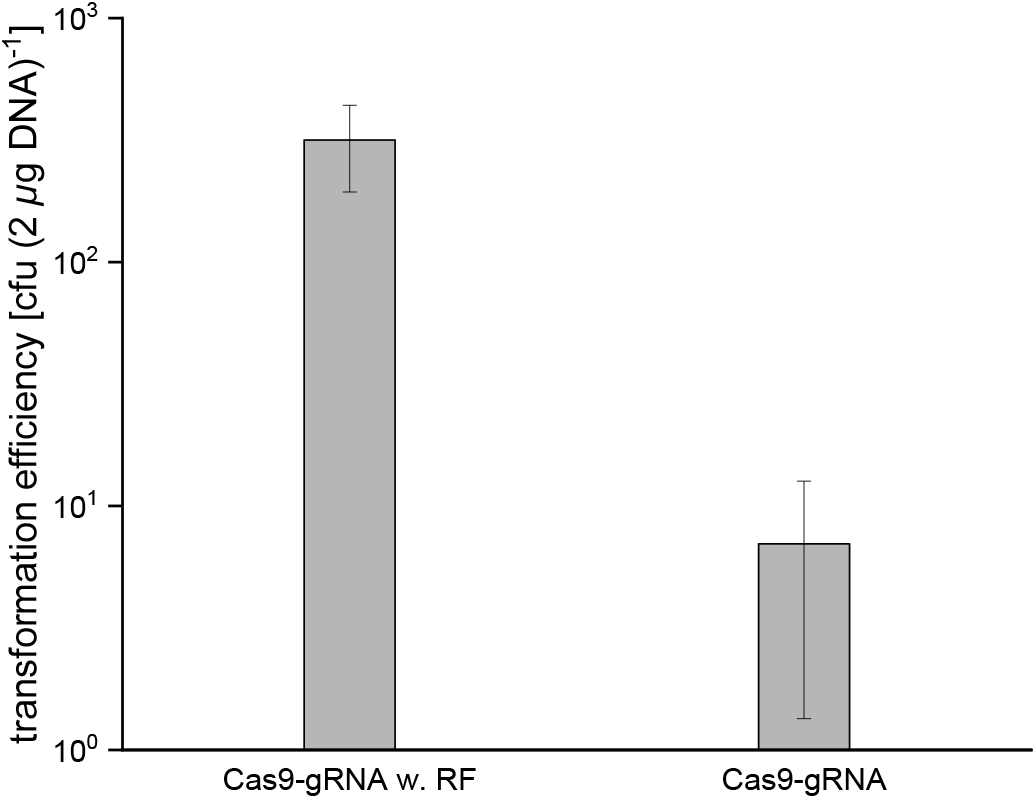
Colony forming units (cfu) for experiments using the CRISPR/Cas9 toolbox. The alanine dehydrogenase-alanine racemase (*ald-alr* 1.9 kbp) genes have been replaced by a 4.2 kbp fragment. The repair fragment (RF) has been inserted via a suicide plasmid. 2 *μg* of Cas9 plasmid and 2 *μg* of the suicide plasmid was co-transformed. The number above the bar represents the positive rate obtained via colony PCR (8 out of 10 were positive). The error bars represent the standard deviation of the biological triplicates.

**Figure S10.**
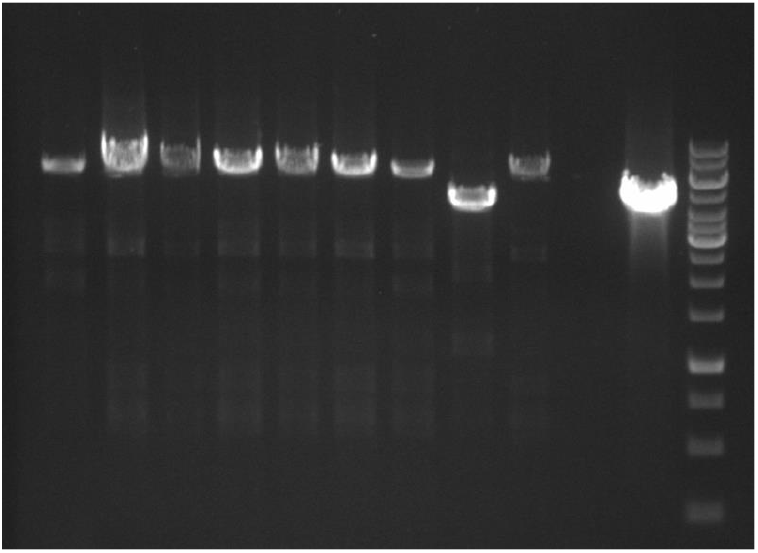
Colony PCR result of CRISPR/Cas9 toolbox edited colonies. Lane 1-10: individual colonies. Lane 11: wildtype. Lane 12: DNA ladder (Thermoscientific GeneRuler 1 kbp DNA ladder #SM0311). One pair of primers was used to check whether the *ald-alr* ORF was replaced with the 4.2 kbp fragment. The expected size of the amplicon of the edited transformants and wildtype genome is 6314 and 4367 bp, respectively.

**Figure S11.**
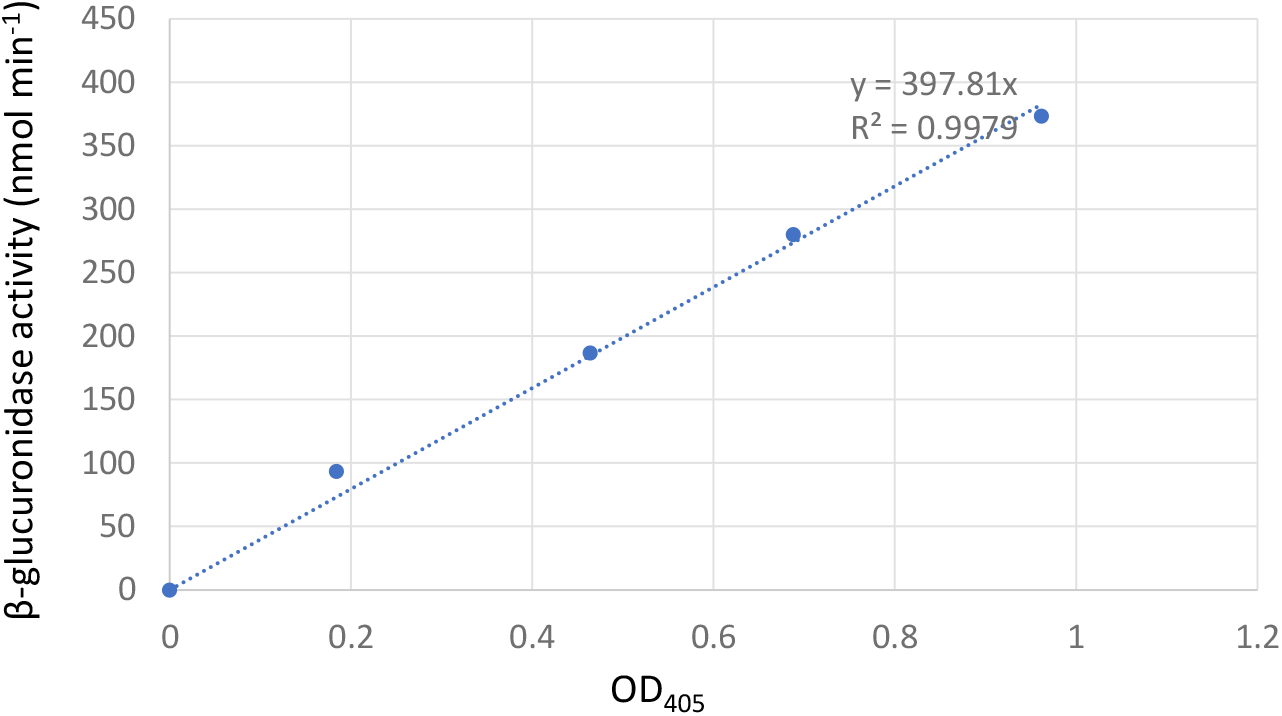
The standard curve of β-glucuronidase activity. Linear regression forcing the curve through point 0/0 yielded the conversion factor 398 nmol min^-1^, that has been applied to all experiments.

